# 3D Structure Determination of Protein Complexes using Matrix-Landing Mass Spectrometry

**DOI:** 10.1101/2021.10.13.464253

**Authors:** Michael S. Westphall, Kenneth W. Lee, Austin Z. Salome, Jean Lodge, Timothy Grant, Joshua J. Coon

**Affiliations:** Department of Biomolecular Chemistry, University of Wisconsin-Madison, Madison, WI; Department of Chemistry, University of Wisconsin-Madison, Madison, WI; Department of Biochemistry, University of Wisconsin-Madison, Madison, WI; Morgridge Institute for Research, Madison, WI

## Abstract

Native mass spectrometry (MS) is an emerging technology that can provide complementary data to electron microscopy (EM) for protein structure characterization. Beyond the ability to provide mass measurements of gas-phase biomolecular ions, MS instruments offer the ability to purify, select, and precisely control the spatial location of these ions. Here we present a modified Orbitrap MS system capable of depositing a native MS ion beam onto EM grids. We further describe use of a chemical landing matrix that both preserves and protects the structural integrity of the deposited particles. With this system we obtained the first 3D reconstructed structure of gas-phase, deposited biomolecular ions – the 800 KDa protein complex GroEL. These data provide direct evidence that non-covalent protein complexes can indeed retain their condensed-phase structures following ionization and vaporization. Finally, we describe how further developments of this technology could pave the way to an integrated MS-EM technology with promise to provide improved cryo-EM sample preparation over conventional plunge-freezing techniques.

---

By allowing so-called elephants to fly, electrospray ionization (ESI) coupled with mass spectrometry (MS) has transformed our ability to characterize proteins.^1^ Perhaps no area better exemplifies this than native MS, where intact protein complexes are gently ionized, vaporized, and mass analyzed.^2^ These mass measurements can provide invaluable information on sub-unit stoichiometry, connectivity, and even the presence of non-covalently bound ligands.

Whether ionized protein complexes truly retain their structure in the gas-phase has been debated for decades.^3^ Collisional cross-sections of numerous gas-phase protein complexes have been experimentally determined by ion mobility MS.^4-6^ In general, these collective data indicate that under optimal conditions the measured cross-sections are consistent with condensed-phase structures. Seeking a direct measurement, Robinson and co-workers configured a quadrupole time-of-flight (q-ToF) MS with a transmission electron microscopy (TEM) grid holder and deposited ions of GroEL and ferritin.^7,8^ These studies imaged particles of roughly the correct size and shape via negative and positive staining TEM. The resultant images, however, lacked the higher resolution features typical of a conventional staining experiment. This lack of detail opens the possibility of structural damage occurring during the experiment and prevents solving the 3D structure from the images. More recently, Longchamp *et al*. imaged soft-landed small proteins using low-energy electron holography followed by numerical reconstruction – again, confirming the ability to soft land onto a surface. However, given the small size of the molecules imaged and lack of details in the reconstructed images it is difficult to tell whether the landed proteins are damaged or not.^9^

Building from this pioneering work, we explored new configurations and approaches for depositing gas-phase protein complexes onto surfaces for direct EM imaging. First, we modified a quadrupole Orbitrap hybrid system (Ultra-High Mass Range Q-Exactive^10^) by removing the collision cell and modifying the end vacuum cap to allow for an insertion probe to hold a TEM grid to the rear of the c-trap exit lens (**Extended Data 1**). With this modification, the ion beam can either be mass analyzed or directed towards a TEM grid.

To test the apparatus, we analyzed the bacterial chaperonin GroEL, an ∼ 800 kDa homo-oligomer having 14 identical subunits, as it has been extremely well-characterized both by native MS and microscopy.^11^ Using nano-electrospray ionization we observed charge states ranging from +71 to +62 across the *m/z* range of 11,000 to 13,000 and having the calculated molecular weight of 802,500 ± 300 Da (**Figure 1A**). Next, we deposited the GroEL ion beam for durations of 60 to 600 seconds onto glow discharged carbon coated TEM grids. After deposition, the TEM grids were removed from vacuum, stained with uranyl acetate, and immediately viewed using TEM (**Figure 1B**). Particles were observed across much of the TEM grids at reasonably high densities; however, the structural features that define GroEL, *i*.*e*., rings and lines, were not observed. Instead, irregular shaped, but mainly featureless particles of approximately the size of GroEL were observed. These images reproduced well those reported by Robinson *et al*.^7,8^

**Figure 1.**
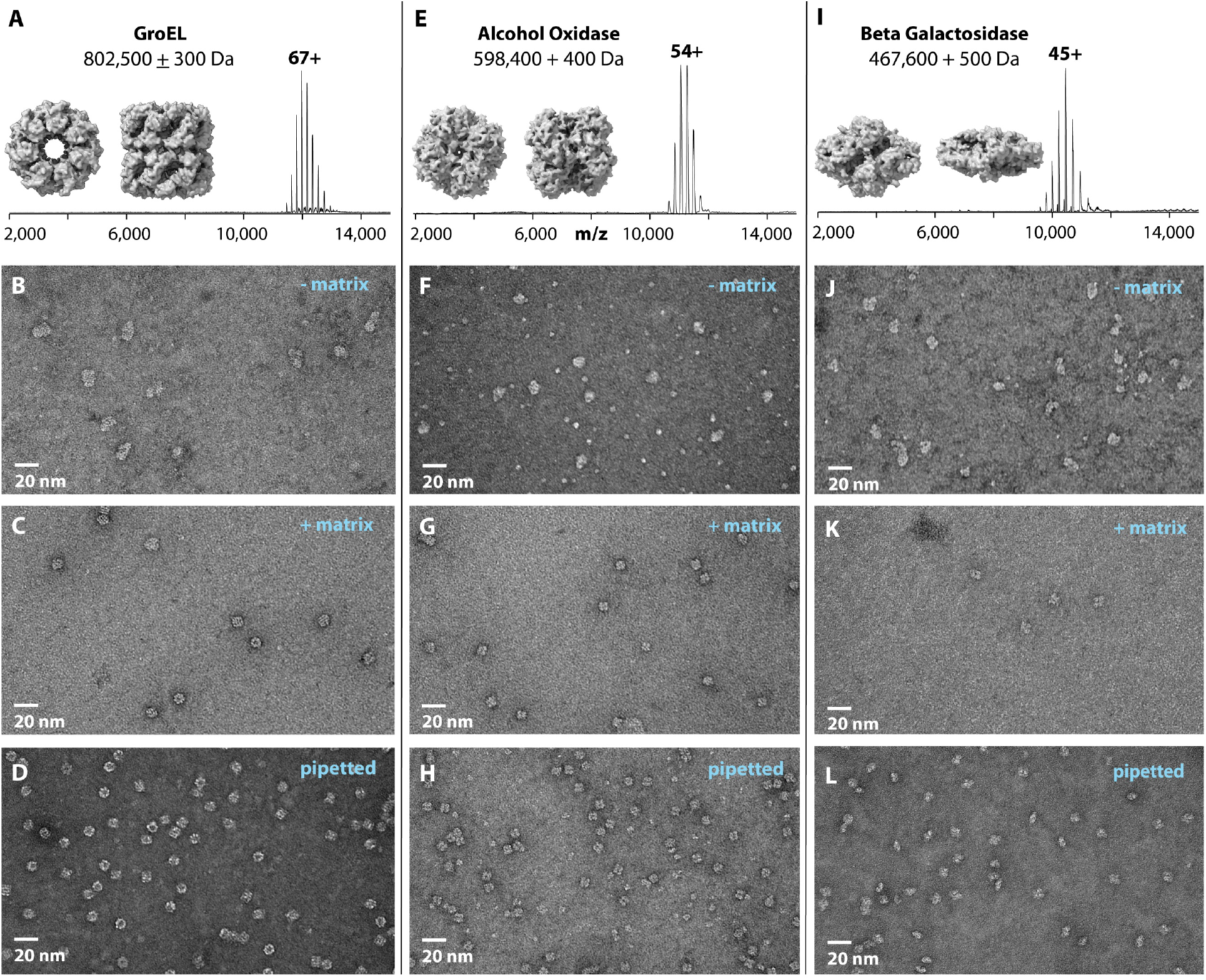
Matrix-landed protein complexes retain structural features and integrity as imaged by negative stain TEM. (A) Native mass spectrum of GroEL complexes along with molecular model images of GroEL. (B) Negative stain TEM image of GroEL ions landed onto bare carbon TEM grids. (C) Negative stain TEM image of GroEL ions matrix-landed onto TEM grids having a thin film of glycerol. (D) Negative stain TEM image of GroEL molecules that were conventionally prepared. (E) Native mass spectrum of alcohol oxidase complexes along with molecular model images of alcohol oxidase. Negative stain TEM images of alcohol oxidase ions landed on bare carbon TEM grids (F) or TEM grids coated with a thin film of glycerol – i.e., matrix-landed (G). (H) Conventionally prepared negative stain TEM images of alcohol oxidase molecules. (I) Native mass spectrum the β-galactosidase complex along with molecular model images of β-galactosidase. Negative stain TEM images of β-galactosidase ions landed onto either bare carbon TEM grids (J) or TEM grids coated with a thin film of glycerol – *i*.*e*., matrix-landed (K). (L) Conventionally prepared negative stain TEM images of β-galactosidase molecules.

From these data we supposed that the GroEL ions had lost their condensed-phase structure via: (1) the process of ionization, vaporization, and transport through MS, (2) lengthy exposure to high vacuum without solvent (*i*.*e*., up to 600 seconds on surface), and/or (3) interactions with the TEM grid surface. To probe this issue, we altered the TEM grid surface by addition of a chemical matrix. Cryoprotective compounds (*e*.*g*., glycerol, trehalose, glucose, ionic liquids, *etc*.) can promote preservation of protein structure, even when dehydrated and/or in vacuum environments; further, several studies have shown the benefits of direct TEM imaging from sugar-fixed particles.^12-15^ Additionally, two MS studies reported using glycerol-coated deposition surfaces.^16,17^ Following these leads, we produced a uniform thin film of glycerol matrix by depositing a small volume (∼ 3 µL) of glycerol onto the carbon TEM grid surface followed by manual blotting, insertion into the modified Orbitrap, and GroEL cation beam deposition. After a deposition period of up to 600 seconds, the TEM grids were removed and negatively stained as described above. With the glycerol film present, similar GroEL particle distributions were observed; however, the particles displayed the characteristic features expected for negatively stained GroEL (**Figure 1C**) as observed in the conventionally prepared sample (**Figure 1D**). Note the clearly visible seven member rings (end views) and the four lines indicating the tetrameric stacked rings (side views) for both the matrix-landed and conventional samples.

To test whether this phenomenon was unique to GroEL, we explored the deposition behavior of two other well-studied protein complexes. The first, alcohol oxidase (AOX, a homo-octamer)^18^ was similarly subjected to nano-electrospray and measured at a mass of 598,400 ± 400 Da (**Figure 1E**). Deposition of these ions onto the bare TEM grids again resulted in visibly distorted particles of varying size (**Figure 1F**). **Figure 1G** displays an image of these same cations when deposited onto glycerol-treated TEM grids. As with GroEL, the expected structural features of the AOX complex are preserved in the presence of the matrix (conventionally prepared AOX shown in **Figure 1H**). Next, we analyzed the tetrameric β-galactosidase complex which comprises four identical 1023 residue long sub-units.^19^ The smallest of the three complexes studied here, β-galactosidase weighs in at just under 500 KDa (**Figure 1I**). Panels J-K of **Figure 1** illustrate the same trend as observed with GroEL and AOX – the glycerol matrix preserves the landed cationic protein complex.

Following our initial supposition, these data provide strong evidence that either extended exposure to vacuum and/or interactions with the TEM grid surface are likely the key factors governing the structural preservation of landed particles. At present, we believe both phenomena are likely at play and, to some extent, are mitigated by the matrix. First, few quality particles are observed following extended vacuum exposures (> 30 minutes) of a matrix-landed protein complex following negative staining, underscoring the importance of a high flux ion beam to reduce the time required for sample preparation. Second, because glycerol is a dielectric, we hypothesize that it may serve to prevent neutralization of the ionic protein complex by insulating it from the conducting carbon surface. When cationic proteins are reduced on surfaces, electron-based dissociation can occur.^20^ Charge neutralization without the stabilizing forces of water could also result in structural deformation. Supporting this idea, attempts to land GroEL cations into an ionic liquid matrix resulted in no observable particles (data not shown). However, this same solution protects and preserves neutral protein complexes from the deleterious effects of vacuum (data not shown). Understanding the mechanisms and roles of the matrix are the subject of ongoing investigations.

Having established a method to deposit and preserve protein complexes from a gaseous ion beam, we sought to probe the decades old question of whether gas-phase biomolecular ions retain their condensed-phase structures. From a GroEL matrix-landed grid we collected a dataset of 340 images on an L120C microscope equipped with a Ceta camera. We expected that with these matrix-landed molecular images we could generate medium resolution (∼ 20 Å) negative stain 3D reconstruction. From the resultant images, having a pixel size of 3.2 Å, we picked and processed 3,500 particles using *cis*TEM.^21^ 2D Classification was performed and ∼2,700 particles contained in the high-quality class averages (**Figure 2A**) were carried forward for further refinement. *Ab-initio* reconstruction and auto-refinement assuming D7 symmetry resulted in the 3D reconstruction shown in **Figure 2B. Figure 2C** displays the superposition of the matrix-landed 3D reconstruction of GroEL (gray) with the high-resolution crystal structure (blue ribbon). These data show excellent agreement between the landed particles and previously determined GroEL structure (PDB:5W0S). As a further control we also obtained a reconstruction of the same sample prepared conventionally (**Extended Data 2**). This comparison also resulted in strong agreement. These results provide the highest resolution experimental data collected to date confirming that non-covalent protein complexes subjected to ionization and mass spectrometry can retain their condensed-phase structures.

**Figure 2.**
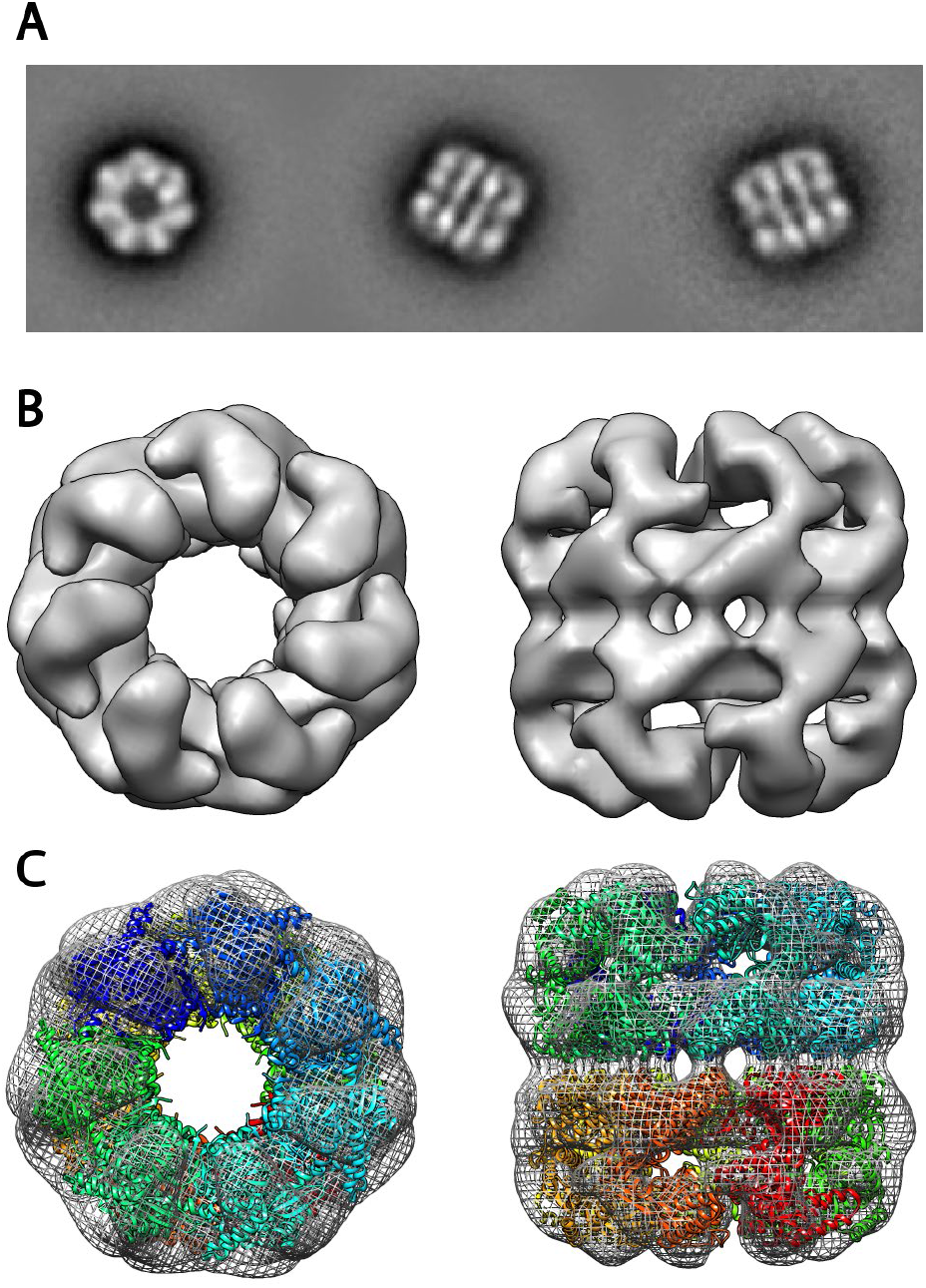
3D reconstruction of GroEL complexes that were matrix-landed from the ion beam of a modified Orbitrap mass spectrometer. (A) 2D Class averages obtained from negatively-stained matrix-landed GroEL cations. (B) Top and bottom views of a three-dimensional reconstruction of GroEL made from the particles contained within the class averages shown in A. (C) Fit of previously determined GroEL structure (PDB:5W0S) into the reconstruction shown in B demonstrating good overall structural agreement.

Here we describe a technique – matrix-landing – that promotes the preservation of structure in non-covalent protein complexes that have traversed a mass spectrometer and been deposited onto a TEM grid. Negative staining provides a simple and robust method to image these structures and further data processing of the negatively stained images can offer detailed structural information. We envision matrix-landing being applied to mixtures of protein complexes where the MS filtering capabilities enable gas-phase purification of the desired particles. Further, a wide variety of native MS technologies exist – *e*.*g*., ion mobility^22^, surface-induced dissociation^23^, collisions^24^, and photo-activation^25^ – and may be utilized to characterize protein complexes and their interactions beyond intact mass measurement. The ability to image the products of these processes could prove invaluable both for structural biology and to the study of gas-phase chemistry.

Detailed evaluations of several relevant native MS conditions (*e*.*g*., buffer, ion source, desolvation energies, time-in-transit, etc.) should also yield further insights and improvements. Exploration of other chemical matrices and methods for generating the highest performing matrix films are similarly critical. Landing conditions likewise play an essential role mandating thorough characterization of ion beam energetics, landing pressures, and effects of vacuum exposure. We suppose that precise control of the TEM grid surface temperature could also be key for maintenance of the ideal matrix surface and for high-resolution structural preservation. Note all the depositions described herein were conducted at room temperature; moving forward we will explore reduced TEM grid temperatures aiming to both extend the protective effects of glycerol beyond ten minutes and to aid in retention of any water/solvent associated with the protein complex ion.

The ultimate extension of this reduced-landing temperature concept is to deposit partially hydrated and mass selected samples directly onto cryogenically-cooled (< 180 ºC) TEM grids. An expected thin coating of amorphous ice would provide protection from the deleterious effects of both vacuum and the TEM grid surface. Directly coupling cryo-EM grid preparation to MS could provide a host of advantages over conventional grid preparation including improved signal and decreased beam-induced motion (due to lower ice background and lack of in-built ice strain^26^, respectively). Aside from the aforementioned benefits derived from MS, the boosted signal could increase resolution and enable imaging of smaller particles. We conclude that the ability to deposit and preserve protein complexes within a vacuum environment may allow for improved TEM sample preparation capabilities and facilitate integration of two disparate fields into one unified technology.

## ACKNOWLEDGEMENTS

We are grateful for support of this project by the National Institutes of Health R35GM118110 grant (to JJC) and especially the program staff for their encouragement, the Morgridge Institute for Research, and the University of Wisconsin-Madison. We thank Mike Sussman, Lloyd Smith, Yifan Cheng, Jim Wells, Paul Ahlquist, Desiree Benefield, Andy Ottens, and Julia Laskin for helpful discussions and the UW-Madison Cryo-EM Research Center for access to TEM instrumentation.

## AUTHOR CONTRIBUTIONS

MSW and JJC conceptually designed the research; MSW constructed the instrumentation; MSW, KWL, AZS, JL, and TG performed the experiments; TG conducted 3D structural reconstructions; MSW, TG, and JJC wrote the paper.

## METHODS

### Materials

Water (Optima LC/MS grade, W6-4) and methanol (Optima for HPLC, A454SK-4) were purchased from Fisher Chemical. Ammonium Acetate (431311-50G), Glycerol (for molecular biology, G5516-100ml), Amicon Ultra-0.5 centrifugal filter (Ultracel-100 regenerated cellulose membrane, UFC510024). Alcohol Oxidase (Pichia pastoris buffered aqueous solution, 55 mg protein/mL, A2404-1KU), GroEl (Chapernion 60 from Escherichia, C7688-1MG), and β-Galactosidase (from Escherichia coli, G3153-5MG) were purchased from Sigma Aldrich. Uranyl Acetate (1% solution, 22400-1) and TEM grids (Carbon support film on 400 mesh copper, CF400-CU) were purchased from Electron Microscopy Sciences.

### Sample preparation

GroEL was prepared at 1 mg/mL in 100 mM ammonium acetate. 200µL of acetone was added to 100µL of buffered protein solution and allowed to sit for five minutes to precipitate the protein. The sample was centrifuged (Fisherbrand Gusto Mini tabletop centrifuge) and the remaining solvent was removed from the pellet. Following this, 400µL of buffer was added to redissolve the protein and placed in an Amicon centrifugal filter. The sample was centrifuged (Thermo Scientific Sorvall Legend Micro 21R) at 10,000 g for 10 minutes at 4°C. Buffer which passed through the filter was discarded and an additional 400µL of buffer was added to the sample for another round of washing using the same centrifugal settings. To obtain the sample, the filter was inverted and centrifuged at 2000 g for 1 minute and diluted with 80µL of buffer.

Alcohol Oxidase was thawed on ice and 10µL was taken and diluted in 390µL of buffer. This solution was buffer exchanged by transfer to an Amicon spin filter and centrifuged at 10,000 g for 10 minutes at 4°C. This cycle was repeated three times replenishing with 400µL of 100mM ammonium acetate after each spin. To obtain the final sample the solution remaining above the filter was removed via pipette.

β-Galactosidase was prepared at 1 mg/mL in 100mM ammonium acetate and buffered exchanged as per Alcohol Oxidase. To obtain the final sample, the filter was inverted and centrifuged at 2000 g for 1 minute and diluted with 200µL of buffer.

### Mass Spectrometry

All mass spectrometry experiments were performed on a modified Thermo Scientific Q-Exactive UHMR Hybrid Quadrupole-Orbitrap mass spectrometer.^10^ Modifications included the removal of the HCD cell ion optics along with the HCD cell vacuum chamber rear cover plate. The cover was replaced with modified plate containing a ball valve assembly which could be evacuated by the roughing pump of the UHMR system. The valve was placed on center with the HCD cell. This arrangement allowed for the use of an insertion probe to place and hold a TEM grid at the exit of the c-trap/entrance to the HCD cell without breaking vacuum on the mass spectrometer. No changes were required to the mass spectrometer electronics or software. Borosilicate glass capillaries were pulled in-house using a model P-2000 laser-based micropipette puller (Sutter Instrument, CA) to an emitter inner diameter of 1 to 5 microns. A platinum wire placed within the capillary provided continuity between the mass spectrometer ESI power supply and the solution being sprayed. All full scan MS1 experiments were conducted with an ESI voltage of 1.1kV to 1.5kV, mass resolving power of 6250 at *m/z* 400, inlet capillary temperature of 250°C, and in-source trapping with -100V offset. For landing experiments, the Obitrap mass analyzer was not employed, and the ions were not stopped within the c-trap. To prevent trapping, the trapping gas pressure was set to a value of 0.1. A decreasing DC gradient was placed on all the ion optics from the inlet of the mass spectrometer to the TEM grid. Specifically, lens voltages of to 20V, 19V, 18V, 17V, 16V, 15V, and 0V were employed on the injection flatapole, inter flatapole lens, bent flatapole, transfer multipole, C-trap entrance lens, and TEM grid respectively. All voltages are adjustable through the user interface with exception of the TEM grid which is tied to ground. No insource trapping was employed, the inlet capillary temperature set to 30°C, and a wide mass filter isolation of 10,000–20,000 *m/z* for GroEL and 8,000–18,000 *m/z* for Alcohol Oxidase and β-Galactosidase. All protein complex solutions were sprayed at a concentration of approximately 0.1 to 0.3 mg/µL.

### Matrix-Landing

Plasma treated carbon film TEM grids were coated with 3 µl of the glycerol/methanol mix (50-50 by volume) and allowed to sit for 30 seconds. Excess solution was removed by touching the edge of the grid to a piece of filter paper. The remaining solution was allowed to equilibrate for 10 minutes. The grid was then placed within the mass spectrometer using the insertion probe and exposed to the ion beam for up to 10 minutes. Upon removal, the grid was negative stained with 75µL of 1% uranyl acetate by edge blotting.

### Transmission electron microscopy

TEM studies of for the reconstruction of negatively stained GroEL were performed at ambient temperature on a Talos L120C (Thermo Fisher Scientific) operated at 120 kV. Electron micrographs were recorded on a Ceta 16 Mpix camera (Thermo Fisher Scientific). All other TEM studies were performed at ambient temperature on a Techai G2 Spirit BioTwin (Thermo Fisher Scientific) fitted with a NanoSprint15 MK-II 15 Mpix camera (AMT Imaging), also operated at 120 kV. For the 3D reconstruction a defocused image series ranging from 0.1 µm to 2 µm in 0.1 µm steps were collected using the SerialEM software package (https://bio3d.colorado.edu/SerialEM/). Particle picking, classification, reconstruction and refinement of 3D maps were performed in cisTEM.

## EXTENDED DATA

**Extended Data 1.**
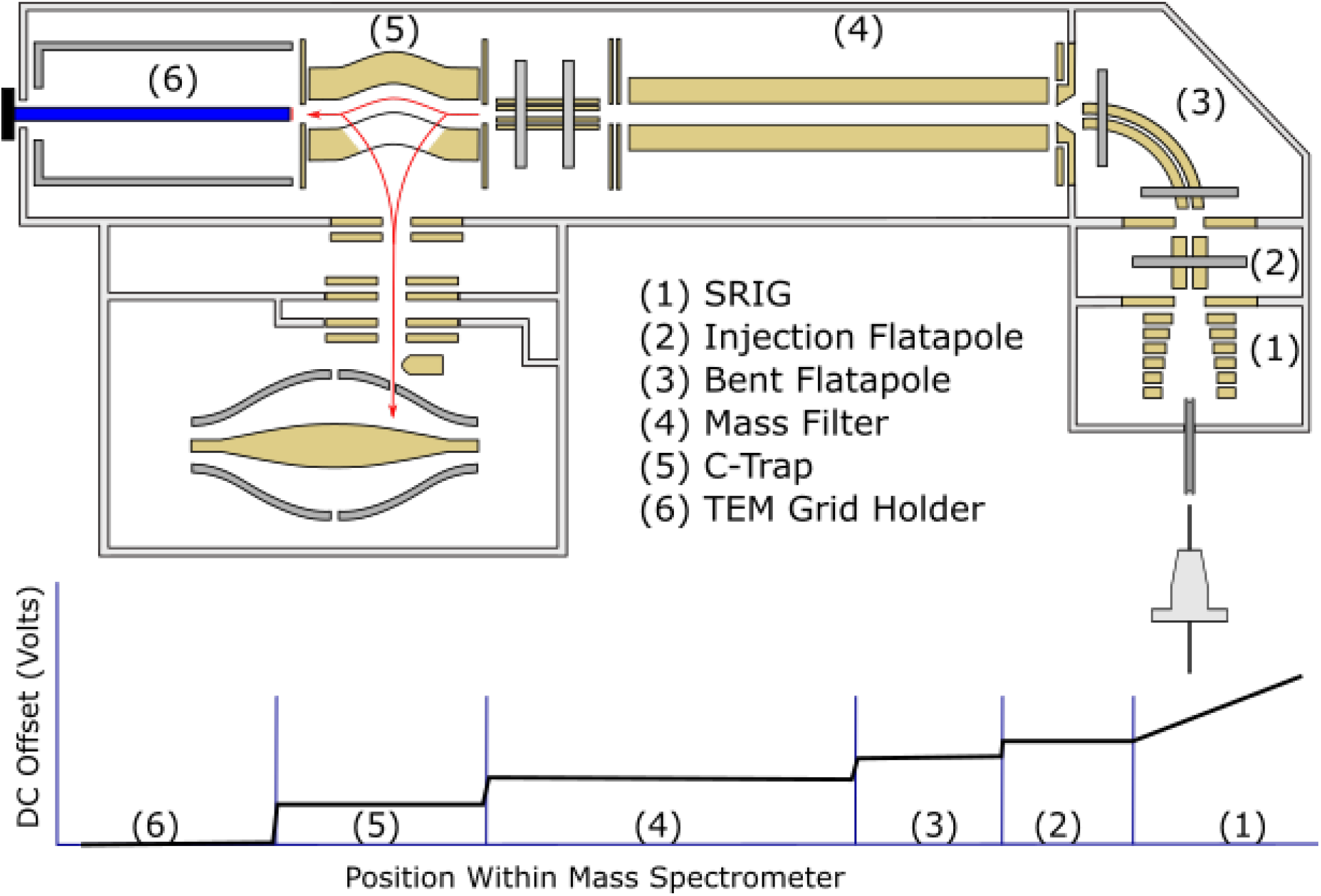
Orbitrap mass spectrometer modified to conduct matrix-landing of biomolecular ions. Ions are generated by nano-electrospray and initially injected into the stacked ring ion guide (SRIG, 1). The ions are then moved through the MS system and either injected into the Orbitrap mass analyzer for mass measurement or directed to the TEM grid (6). The DC offset potentials used during landing experiments is shown on the bottom panel.

**Extended Data 2.**
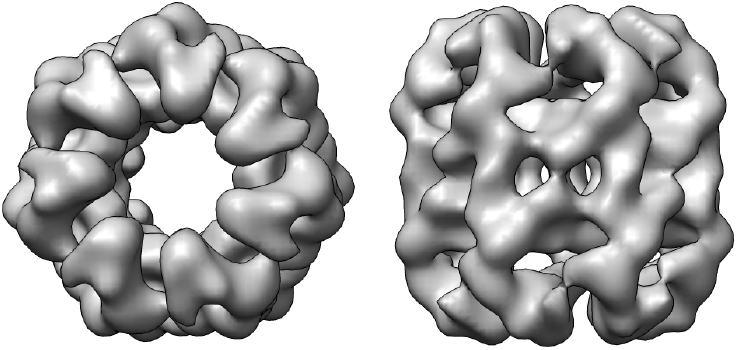
Top and bottom views of a three-dimensional reconstruction of GroEL calculated from images of conventionally prepared negative-stain grids. The images were processed using the same processing procedure as for the matrix-landed reconstruction shown in Figure 2 and the reconstruction contains approximately the same number of particles.

